# Deletion of the Creatine Transporter in dopaminergic neurons leads to hyperactivity in mice

**DOI:** 10.1101/293563

**Authors:** Zuhair I. Abdulla, Bahar Pahlevani, Jordan L. Pennington, Nikita Latushka, Matthew R. Skelton

**Affiliations:** Division of Neurology, Cincinnati Children’s Research Foundation, Cincinnati, OH, USA and Department of Pediatrics, University of Cincinnati College of Medicine, Cincinnati, OH, USA

**Keywords:** Creatine, creatine transporter, Parkinson’s disease, DAT-Cre, hyperactivity, motor function

## Abstract

Creatine (Cr) is required for proper neuronal function, as evidenced by the intellectual disability and epileptic phenotype seen in patients with cerebral Cr deficiency syndromes (CCDS). In addition, attention-deficit hyperactivity disorder (ADHD) is a frequent co-morbidity of Cr transporter (Crt) deficiency, the leading cause of CCDS. While the effects of the loss of Cr in the whole brain is clear, it is unknown if Cr is required for the proper function of all neurons. Of particular interest are dopaminergic neurons, as many CCDS patients have ADHD and Cr has been implicated in dopamine-associated neurodegenerative disorders, such as Parkinson’s and Huntington’s diseases. The purpose of this study was to examine the effect of a loss of the *Slc6a8* (Cr transporter; Crt) gene in cells expressing the dopamine transporter (*Slc6a3*; DAT) on activity levels and motor function as the animals age. DAT-specific Crt-knockout (DAT-Crt) mice were tested along with control (Crt-FLOX) mice monthly from 3 to 12 months of age in locomotor activity, the challenging beam test, and spontaneous activity. DAT-Crt mice were hyperactive compared with controls and this finding persisted throughout the lifetime of the mice. No changes were observed in errors to cross a narrow bridge in the challenging beam test. In a measurement of spontaneous activity, DAT-Crt mice showed increased rearing and hind limb steps, suggesting the hyperactivity carried over to this task. Taken together, these data suggest that the lack of Cr in dopaminergic neurons causes hyperactivity while sparing motor function.

**Abbreviations:**

Cr: Creatine
CK: Creatine Kinase
P-Cr: Phosphocreatine
PD: Parkinson’s Disease
MPTP: 1-methyl-4-phenyl-1,2,3,6-tetrahydropyridine
6-OHDA: 6-hydroxydopamine
Crt: Creatine Transporter
Crt^-/y^: ubiquitous creatine transporter knockout mouse
DAT-Crt^-/y^: dopamine-specific creatine transporter knockout mouse ADHD: Attention-deficit hyperactivity disorder

## Introduction

Creatine (Cr) is a nitrogenous acid involved in energy homeostasis. Through a Cr kinase (CK)-mediated process, Cr is phosphorylated at sites of energy production and diffuses through the cytosol, creating phospho-Cr (P-Cr) pools (Wallimann et al., 2011; Lowe et al., 2013). At sites of ATP consumption CK transfers the phosphate from P-Cr to ADP (Wyss and Kaddurah-Daouk, 2000; Wallimann et al., 2011). The Cr/P-Cr shuttle is a spatial and temporal ATP buffer, preventing transient changes in ATP:ADP or ATP:AMP ratios from increasing mitochondrial production (Wallimann et al., 2011). High Cr concentrations are seen in the brain, heart, and muscle, reflecting the importance of Cr in energetically-demanding tissue. In addition, Cr has antioxidant, neuroprotective, and mitochondria-protective capabilities (Wyss and Kaddurah-Daouk, 2000). The significance of brain Cr is highlighted by the hallmark phenotypes of intellectual disability and epilepsy in Cr deficiency syndromes (Cecil et al., 2001; DeGrauw et al., 2002; van de Kamp et al., 2013). Further, the involvement of Cr has been implicated in several neurodegenerative diseases, such as Parkinson’s disease (PD) and Huntington’s disease, suggesting that Cr may play an important role in motor function.

PD is a neurodegenerative disorder caused by degradation of dopaminergic neurons projecting from the substantia nigra. While the etiology of PD is unknown, there is a strong association between mitochondrial dysfunction and PD. Many PD-related proteins such as alpha-synuclein are involved in mitochondrial function (Shavali et al., 2008; Bose and Beal, 2016). Further, the mitochondrial toxicants rotenone and 1-methyl-4-phenyl-1,2,3,6-tetrahydropyridine (MPTP) are used in rodents to model the motor impairments related to PD. Since Cr is an important mediator between mitochondrial production and energy consumption, perturbations in Cr may also play a role in the pathophysiology of PD. Cr supplementation prevented motor deficits in MPTP-treated mice (Klivenyi et al., 2004), suggesting that Cr could mediate mitochondrial-induced motor impairment. Additionally, Cr protected dopamine neurons from 6-hydroxydopamine (6-OHDA) induced toxicity (Cunha et al., 2014). Based on the neuroprotective effects of Cr and the relative safety of the compound, a clinical trial using Cr supplementation in PD patients was conducted. While initial reports showed that Cr supplementation improved mood and slowed the escalation of dopaminergic agonist doses (Bender et al., 2006), the larger trial showed no improvement in motor symptoms following Cr supplementation (Mo et al., 2017). While the failure of the clinical trials suggest Cr does not improve the motor symptoms of PD patients, it does not exclude the possibility that Cr plays a role in PD. The etiology of PD involves a significant, progressive degradation of dopaminergic neurons and PD symptoms do not manifest until 70-80% of dopamine neurons are lost (Goldberg et al., 2003; Khaindrava et al., 2011). It is possible that the critical window for Cr supplementation, and the involvement of Cr in the motor deficits precedes the manifestation of symptoms. It is also possible that Cr plays an important role in the non-motor symptoms of PD. For instance, mild cognitive impairment is one of the primary non-motor complications of PD (Li et al., 2015). While neither coenzyme Q10 nor Cr alone are able to ameliorate the cognitive decline associated with PD, a recent trial has shown that combining the two compounds is effective (Li et al., 2015), suggesting an adjunctive role for Cr in the treatment of PD.

To better understand the role of brain Cr, we developed transgenic mice with exons 2-4 of the murine *Slc6a8* (creatine transporter; Crt) gene flanked by loxP sites (*Slc6a8^Flox^*; Skelton et al., 2011). The *Slc6a8* gene is responsible for the transport of Cr into cells (Wyss and Kaddurah-Daouk, 2000). Ubiquitous knockout (*Crt^-/y^*) mice generated from this line have reduced cerebral Cr, lower body weight, and show significant learning and memory deficits (Skelton et al., 2011; Hautman et al., 2014). Additionally, *Crt^-/y^* mice have reduced swim speed compared with control (*Crt^+/y^)* mice, suggesting that these mice may have motor deficits (Skelton et al., 2011). Isolated hippocampal mitochondria from *Crt^-/y^* mice exhibit increased respiration and had higher mitochondrial content, but an overall decrease in cerebral ATP levels (Perna et al., 2016). In order to isolate *Slc6a8*’s influence on the dopaminergic system, and to reduce potential confounds created by the size differences in ubiquitous knockout mice, we created a dopamine-specific *Slc6a8* knockout mouse (DAT-Crt^-/y^). The purpose of this study was to determine if mice carrying a dopamine-specific knockout of *Slc6a8* would show motor deficits. The main finding of this study was that deleting the Crt from the dopaminergic system in mice results in hyperactivity.

## 2. Experimental Procedures

### 2.1 Animals

Slc6a3^tm1.1(cre)Bkmn^/J (DAT^Cre^) mice were obtained from Jackson Labs (Bar Harbor, ME) and bred to wild-type mice from the same background strain (C57Bl6/J) for at least one generation to establish the in-house colony. In-house generated female Slc6a8^fl/fl^ mice (Skelton et al., 2011) were mated with male DAT^Cre^ mice. The resulting offspring were used as subjects for these studies with no more than 1 animal/genotype/litter represented. Male Slc6a8^fl/y^::DAT^Cre+^ (DAT-Crt^-/y^) and Slc6a8^fl/y^::DAT^Cre-^ (FLOX) were tested every 30 days beginning at postnatal day (P) 90 and ending on P360. We have shown that the behavior of the FLOX mice does not differ from WT mice (Udobi et al., 2018). Mice were weighed monthly prior to running behavioral assessments. Thirteen Flox and 11 DAT-Crt^-/y^ mice were enrolled in the study, however several animals died and the study concluded with 8 flox and 7 DAT-Crt^-/y^ animals. All procedures were performed in the Cincinnati Children’s Research Foundation vivarium which is fully accredited by AAALAC International and protocols were approved by the Institutional Animal Use and Care Committee. Lights were on a 14:10 light:dark cycle, room temperature was maintained at 19±1º C. Food and water were sterilized and available *ad libitum.* All institutional and national guidelines for the care and use of laboratory animals were followed.

### 2.2 Behavior

#### 2.2.1 Challenging Beam Task

The protocol was adapted from Fleming et al. (2004) with minor changes. A 1 m beam was divided into four-25 cm segments, the first was 3.5 cm wide and the width of each subsequent segment was reduced in 1 cm increments to a final width of 0.5 cm. A wire mesh grid (0.5 cm x 0.5 cm) was placed atop the beam. The bridge was elevated 13 cm from the tabletop using standard barrier mouse cages. Mice were trained for two days prior to testing. On day one, mice were assisted as they traversed the 1 m beam to the home-cage goal for two trials. Day one concluded when mice were able to complete 5 unassisted bridge traversals. On the second day of training, mice were required to complete 5 unassisted bridge traversals. For testing, mice were assessed over three, 2 min trials. Trials were recorded and scored by a trained observer blinded to genotype. The variables measured were latency to cross (2 min maximum), errors – defined as a foot-slip during forward progress – committed per steps taken, and total number of steps (Fleming et al., 2004).

#### 2.2.2 Spontaneous Activity

A transparent glass cylinder (15.5 cm [h] × 12.7 cm [dia]) was placed on a plexiglass platform with a camera below to record activity. Mice were placed in the cylinder for 3 min. Trials were recorded and scored offline by a trained observer blinded to genotype. The dependent measures were steps per limb (front and hind limb), rears, and total time grooming.

#### 2.2.3 Overnight Locomotor Activity

Locomotor activity was assessed in automated activity monitoring chambers [40 cm (w) × 40 cm (d) × 38 cm (h)] with 16 LED-photodetector beams in the X and Y planes (Photobeam Activity System – Open Field, San Diego Instruments, San Diego, CA). Photocells were spaced 2.5 cm apart. Mice were tested for 14 h, consisting of the last 2 h of the light phase, the 10 h dark phase and 2 h of the following light phase.

#### 2.3 Statistics

Repeated-measures two-way ANOVAs (age x genotype) were used for all analyses in this study. Significant interactions were evaluated using post hoc analysis with the Holm-Sidak correction. Significance was set at p<0.05 for all tests.

### 3. Results

#### Weight

There was an age × gene interaction observed (F(9,153)=1.95, P<0.05) (Fig. 1). DAT-Crt^-/y^ mice weighed less than FLOX mice from P270 to P330. All animals gained weight as they aged (F(9,153)=18.92, P<0.0001).

#### Challenging Beam

For number of steps to cross the beam, there was an age × gene interaction (F(8,104)=4.24, P<0.001; Fig. 2). On P90, DAT-Crt^-/y^ mice took fewer steps to cross the beam than FLOX mice (P<0.001). For number of errors committed there was a significant age × gene interaction (F(8,136)=2.20, P<0.01); however, simple-effect ANOVAs showed no significant effects. For latency to cross the bridge, there were main effects of gene (F(1,17)=5.27, P<0.05) and age (F(8,136)=6.43, P<0.001). DAT-Crt^-/y^ mice traversed the beam faster than FLOX animals and the latency to cross decreased as mice aged.

#### Spontaneous Activity

There was a main effect of gene on rearing behavior with DAT-Crt^-/y^ mice rearing more than FLOX mice (F(1,14)=8.899, P<0.01) (Fig. 3). For number of hind limb steps, there was a main effect of gene (F(1,14)=11.8, P<0.01) with DAT-Crt^-/y^ mice taking more steps than FLOX mice. For grooming duration and forelimb steps, there were no significant effects of gene. There was a main effect of age for forelimb steps with older mice taking less forelimb steps (F(8,112)=3.465, P<0.05).

#### Overnight Locomotor Activity

There were significant main effects of age (F(9,144)=3.89, P<0.001) and gene (F(1,16)=21.09, P<0.001) along with a significant age × gene interaction for total activity (F(9,144)=3.88, P<0.001; Fig. 4). DAT-Crt^-/y^ mice had increased activity compared with FLOX mice at all ages except P330. During the light phase, there were significant main effects of gene (F(1,16)=35.51, P<0.0001), age (F(9,144)=5.05, P<0.0001), and an age × gene interaction (F(9,144)=4.78, P<0.0001) with DAT-Crt^-/y^ mice more active than FLOX mice on at all ages except P330 (Table 1, Fig. 5). Similarly, during the dark phase there were significant main effects of gene (F(1,16)=15.42, P<0.01), age (F(9,144)=2.56, P<0.01), and a significant age × gene interaction (F(9,144)=3.49, P<0.001). As with the other measures, with DAT-Crt^-/y^ mice were more active than FLOX mice on all days tested except P330.

## 4. Discussion

DAT-Crt^-/y^ mice traversed the challenging beam faster than controls, exhibited more stepping activity in spontaneous activity, and displayed hyperactivity in overnight locomotor assessments. These results were consistent across the duration of this experiment, suggesting that deletion of Slc6a8 in the dopaminergic system results in hyperactivity in mice. Our results suggest that the lack of the Cr/P-Cr shuttle plays an important role in dopamine-mediated locomotion.

**Figure 1.**
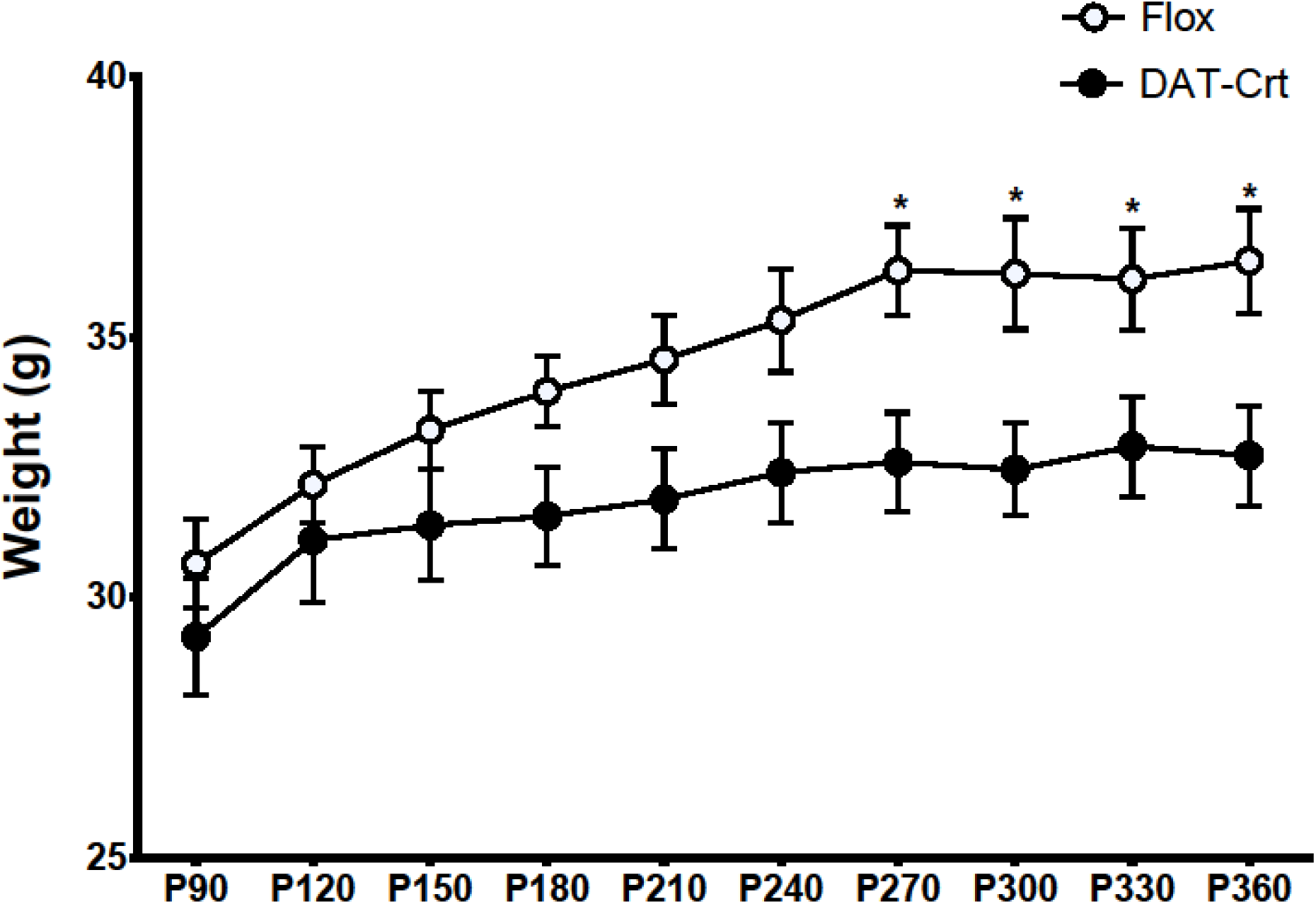
Weight. An age × gene interaction revealed that DAT-Crt^-/y^ mice weighed less than FLOX mice from P270 to P330. All animals gained weight as they aged.

**Figure 2.**
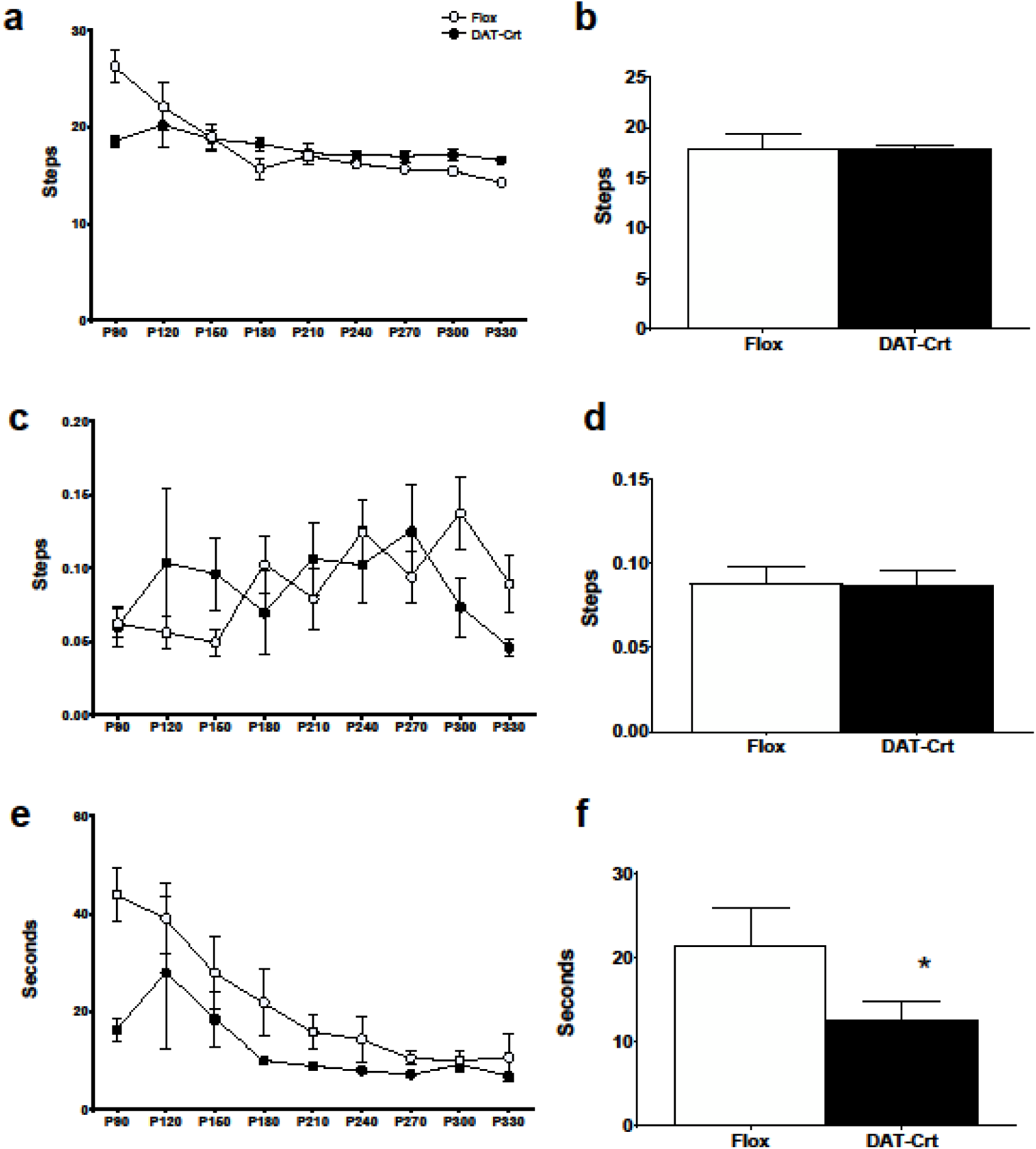
Challenging Beam Task. An age × gene interaction was reported for number of steps required to cross the beam (a, b), with DAT-Crt^-/y^ mice requiring less steps on P90. Although there was a significant age × gene interaction regarding errors (c, d), simple-effect ANOVAs showed no significant effects. Additionally, DAT-Crt^-/y^ mice traversed the beam faster than FLOX animals and the latency to cross decreased as mice aged (e, f).

**Figure 3.**
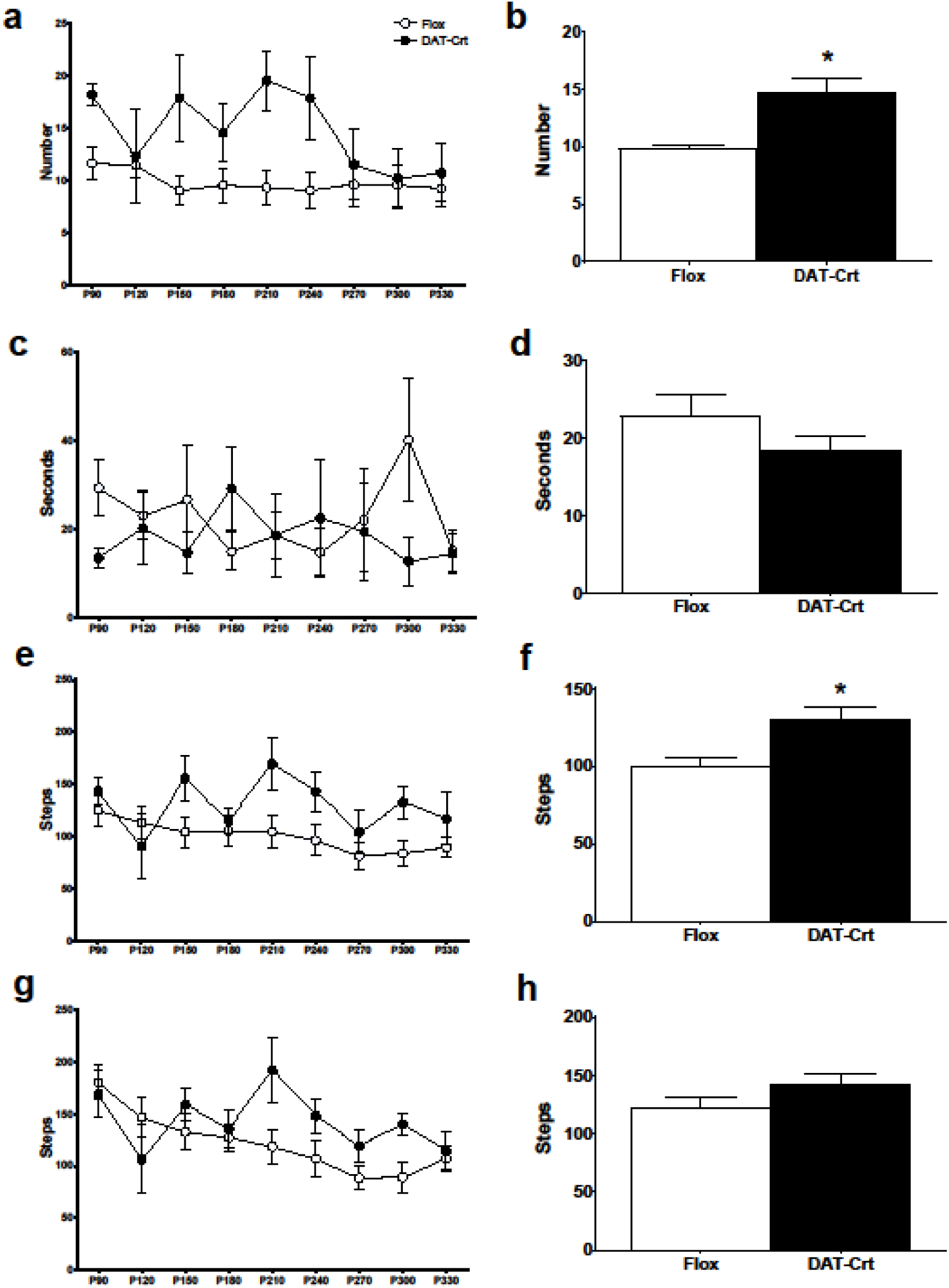
Spontaneous Activity. DAT-Crt^-/y^ mice reared more (a, b) and took more hindlind steps than FLOX mice (e, f), but activity did not differ on grooming duration (c, d) or on forelimb steps (g, h).

**Figure 4/5.**
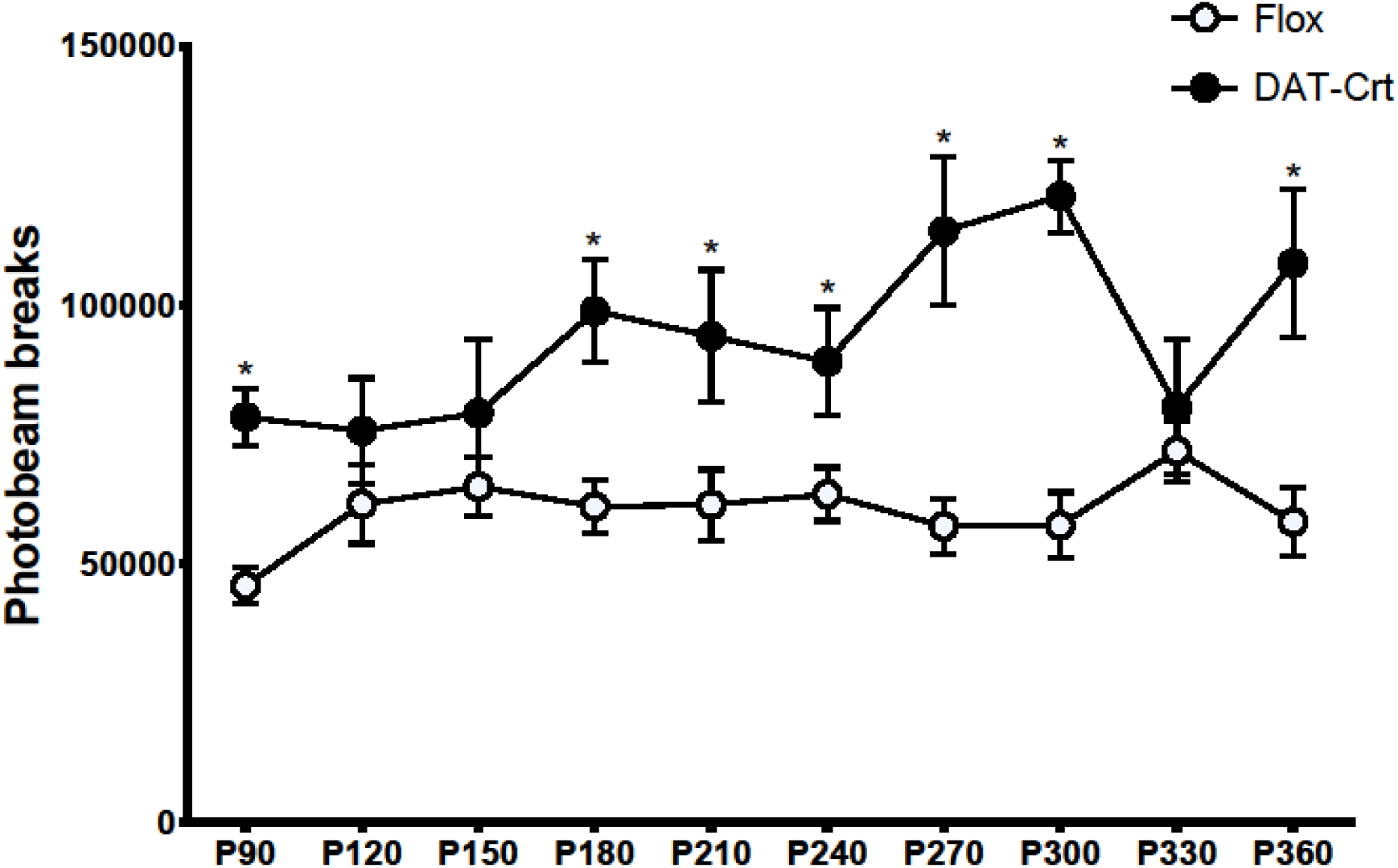
Overnight Locomotor Activity. DAT-Crt^-/y^ mice had increased activity compared with FLOX mice at all ages except P330.

**Figure 5.**
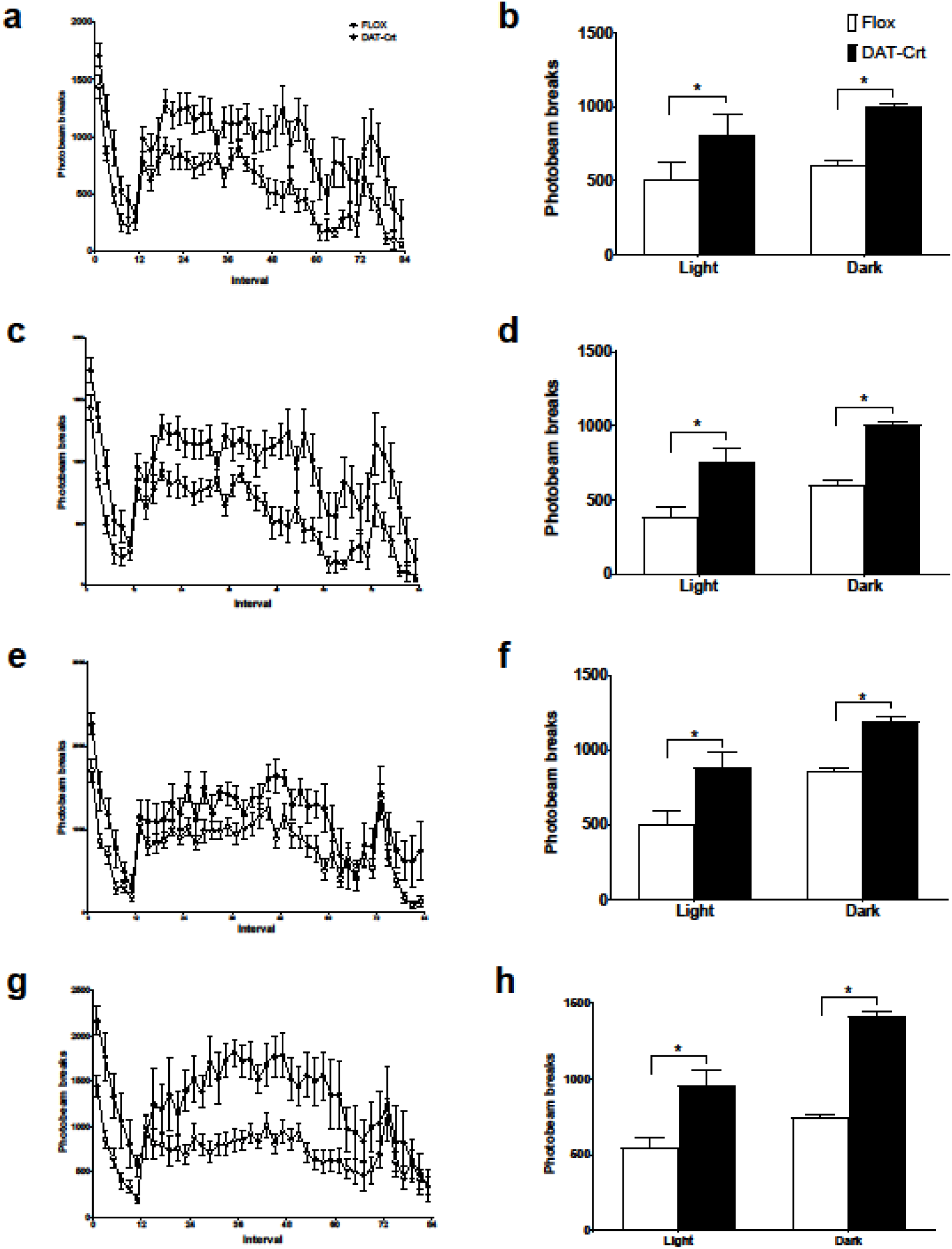
Activity levels during the light and dark phases. Both total activity and light vs. dark analyses revealed that DAT-Crt^-/y^ mice were more active than flox controls on all testing periods except P330. Above, representive graphs for total activity and light vs. dark for P90 (a,b), P120 (c,d), P240 (e,f), and P360 (g,h) mice.

The challenging beam task was designed to uncover subtle motor deficits that could provide translational value to human neurodegenerative disorders (Fleming et al., 2004). Animals exhibiting a PD-like phenotype display increased errors, steps to cross, and latency to cross the beam in this task (Fleming et al., 2004; Glajch et al., 2012). DAT-Crt^-/y^ mice did not show any deficits on challenging beam task. In contrast, DAT-Crt^-/y^ mice actually had shorter latencies when crossing the tapered beam and never relied on more steps to cross the beam than control mice. Coupled with a lack of errors on this task, DAT-Crt^-/y^ performance on the challenging beam task appear to be mediated by a hyperactivity that apparently does not impair motor control.

Furthermore, in the spontaneous activity task, DAT-Crt^-/y^ mice had increased rearing behavior and hind limb steps compared with FLOX mice. Mice with motor deficits typically display a decrease in these behaviors (Schallert et al., 2000; Fleming et al., 2004; Glajch et al., 2012; Pinto et al., 2016). These increases are indicative of hyperactivity (O’Neill and Gu, 2013; Kostrzewa et al., 2015). Similarly, DAT-Crt^-/y^ mice were hyperactive in overnight locomotor activity assessments during both light and dark phases, compared with FLOX controls. This hyperactivity persisted throughout the year-long study, except at P330. Taken together these behavioral measures suggest that the DAT-Crt^-/y^ mice are hyperactive.

The purpose of this study was to determine if DAT-Crt^-/y^ mice show motor deficits that have been used to model a parkinsonian phenotype. The results suggest that DAT-Crt^-/y^ mice do not have motor deficits but are hyperactive. Interestingly, ADHD is frequently comorbid with CTD (van de Kamp et al., 2013). Additionally, 6-OHDA is a toxicant that preferentially targets dopamine neurons (Gainetdinov et al., 1998) and while 6-OHDA is often used to model PD in adult rodents (Schallert et al., 2000; Tirmenstein et al., 2005; Glajch et al., 2012; Cunha et al., 2013, 2014; Kupsch et al., 2014), neonatal administration causes hyperactivity in rats (Kostrzewa et al., 2015). This suggests that certain dopamine-mediated behaviors are governed by critical periods. Cells lacking *Slc6a8* have an altered energetic profile: mitochondrial content is increased, but the mitochondria may be dysfunctional. At a whole-body level, *Slc6a8^-/y^* mice have reduced brain ATP levels and whole-body energy expenditure is increased (Perna et al., 2016). Because several aspects of dopamine release and reuptake require ATP, the loss of P-Cr pools in these neurons appear to disrupt the dopaminergic system (Sulzer et al., 2016). Given the high rate of ADHD in the CTD population, future studies will look for attentional abnormalities in DAT-Crt^-/y^ mice.

Although we did not observe the typical motor symptoms seen in parkinsonian animals, this study did not necessarily confirm a lack of PD-like behavior in DAT-Crt^-/y^ mice. Some estimate that by the time of a diagnosis of PD there is already a 70-80% loss of the neurons in the substantia nigra (Goldberg et al., 2003; Khaindrava et al., 2011). For this reason, it appears that only evaluating DAT-Crt^-/y^ mice for one year may be a limitation to this study. It is not uncommon for animals models of PD to display an initial hyperactivity followed by hypoactivity coupled with parkinsonian symptoms as they age (Havrda et al., 2013; Schultheis et al., 2013). For instance, Atp13a2-deficient mice do not differ from wild type animals on motor tests until 20 months of age (Schultheis et al., 2013). Our animals were only tested for one year, and although these assessments do not point to a PD-like animal, it is possible that trials ended before a PD-like phenotype emerged. Future studies should assess DAT-Crt^-/y^ mice at an older age. Alternatively, because DAT-Crt^-/y^ mice display a robust hyperactivity, this may indicate the presence of an ADHD-like phenotype and potentially explain the high comorbidity of ADHD and CTD.

## Acknowledgments

The authors would like to thank Marla K. Perna, Kenea C. Udobi, and Keila N. Miles for providing scientific input and proofreading this article.

## Author contributions

Mr. Abdulla and Dr. Skelton designed the experiment, drafted and edited the manuscript. Mr. Abdulla, Ms. Pahlevani, Ms. Pennington, and Mr. Latushka conducted and scored all experiments.

